# Contributions of hyperactive mutations in M^pro^ from SARS-CoV-2 to drug resistance

**DOI:** 10.1101/2023.09.28.560010

**Authors:** Julia M. Flynn, Sarah N. Zvornicanin, Ala M. Shaqra, Nese Kurt Yilmaz, Stephanie Moquin, Dustin Dovala, Celia S. Schiffer, Daniel N.A. Bolon

## Abstract

The appearance and spread of mutations that cause drug resistance in rapidly evolving diseases, including infections by SARS-CoV-2 virus, are major concerns for human health. Many drugs target enzymes, and resistance-conferring mutations impact inhibitor binding and/or enzyme activity. Nirmatrelvir, the most widely used inhibitor currently used to treat SARS-CoV-2 infections, targets the main protease (M^pro^) preventing it from processing the viral polyprotein into active subunits. Our previous work systematically analyzed resistance mutations in M^pro^ that reduce binding to inhibitors; here we investigate mutations that affect enzyme function. Hyperactive mutations that increase M^pro^ activity can contribute to drug resistance but had not been thoroughly studied. To explore how hyperactive mutations contribute to resistance, we comprehensively assessed how all possible individual mutations in M^pro^ affect enzyme function using a mutational scanning approach with a FRET-based yeast readout. We identified hundreds of mutations that significantly increased M^pro^ activity. Hyperactive mutations occurred both proximal and distal to the active site, consistent with protein stability and/or dynamics impacting activity. Hyperactive mutations were observed three times more than mutations which reduced apparent binding to nirmatrelvir in recent studies of laboratory grown viruses selected for drug resistance. Hyperactive mutations were also about three times more prevalent than nirmatrelvir-binding mutations in sequenced isolates from circulating SARS-CoV-2. Our findings indicate that hyperactive mutations are likely to contribute to the natural evolution of drug resistance in M^pro^ and provide a comprehensive list for future surveillance efforts.

## Introduction

The emergence of drug resistance in rapidly evolving diseases including cancers and infections leads to poor health outcomes with large costs to human communities^1^. The development of improved treatments has been outpaced by the rate of resistance evolution and if current trends continue drug-resistant infectious diseases will cause 10 million deaths worldwide by 2050^2,3^. Improved strategies to understand and reduce the impacts of drug resistance are urgently needed.

The COVID-19 pandemic, caused by the highly contagious SARS-CoV-2 virus, has had a substantial impact on human health, highlighting the need to develop effective therapeutics with reduced potential for resistance. The SARS-CoV-2 genome encodes two overlapping polyproteins – pp1a and pp1ab^4^. M^pro^, a dimeric cysteine protease encoded by the nsp5 gene, initiates autolytic cleavage out of the polyproteins and then cleaves at an additional 11 conserved sites, releasing the functional proteins required for viral replication and transcription^5–7^. M^pro^ is thus essential in the viral life cycle, making it an attractive target for the design of anti-viral drugs.

Essential enzymes are one of the most common targets of drugs used to treat rapidly evolving diseases^8^; they are often structured, well studied, and amenable to biochemical assays which can drive medicinal chemistry optimization. In the evolution of drug resistance, enzyme targets frequently accumulate mutations that impact the binding of drug and/or the turnover of substrate^9^. The impact of mutations on inhibitor binding are relatively straightforward to interpret and we have reported a comprehensive assessment of how all possible point mutations in M^pro^ from SARS-CoV-2 impact binding to drugs^10^. In addition to drug binding, the effects of mutations on enzyme activity are also critical to understanding drug resistance.

The impacts of mutations on enzyme activity are important for at least two reasons (Figure 1). First, enzyme activity must be high enough to enable virus propagation such that only mutations or combinations of mutations that maintain sufficient activity are accessible for resistance evolution in viral populations. Hyperactive mutations that increase activity beyond that of the wild-type (WT) enzyme can provide an activity buffer that either rescues or enables the evolution of additional mutations (Figure 1A&B). For example, in response to inhibitors of HIV protease, primary mutations that strongly disrupt drug binding tend to also reduce substrate processing and require compensatory mutations that rescue activity for the evolution of high-level resistance^11–13^.

**Figure 1.**
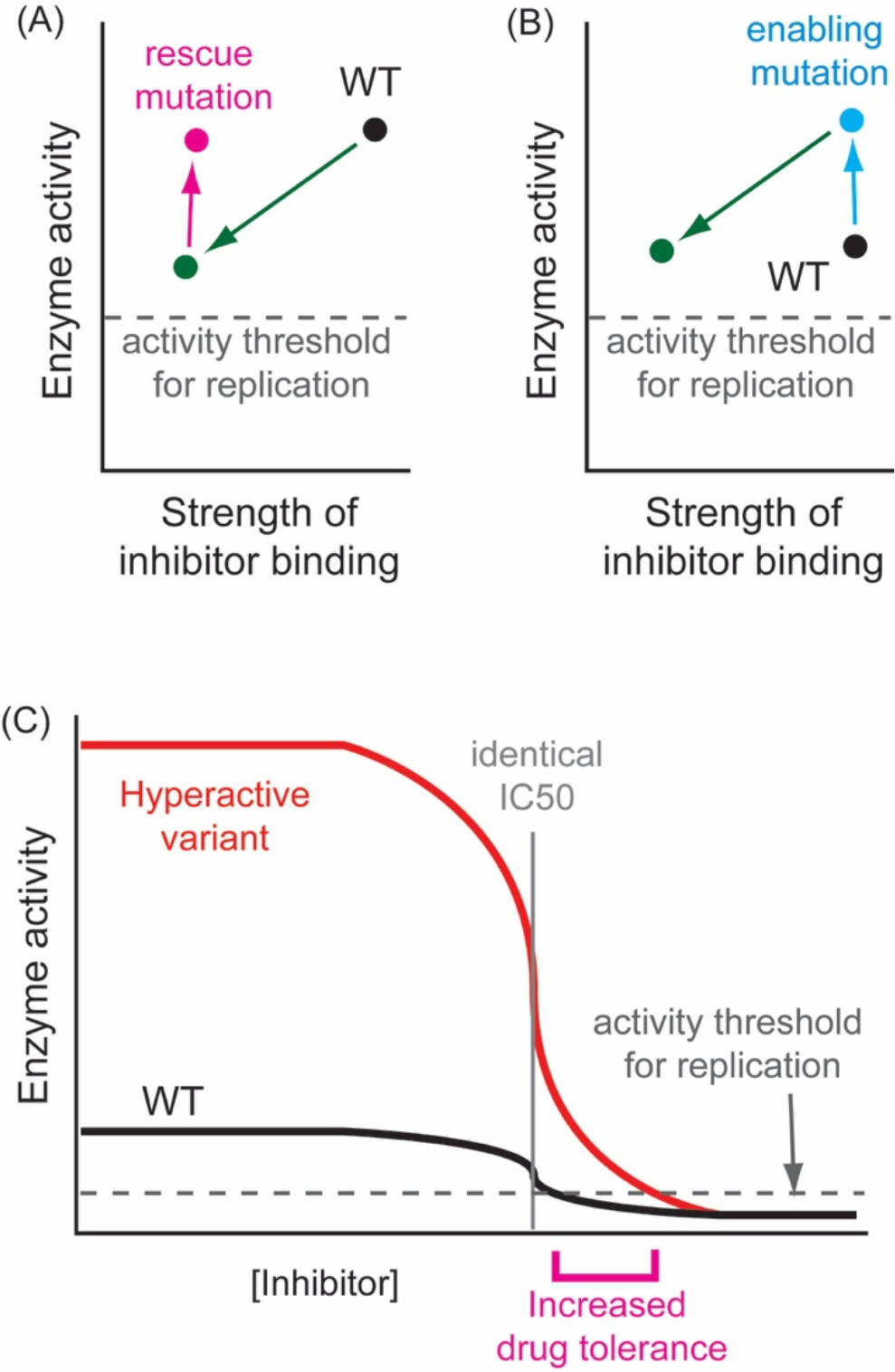
Different mechanistic contributions of hyperactive mutations in the evolution of drug resistance. (A) Rescue mutations elevate enzyme activity back towards WT levels after mutations that both disrupt drug binding and decrease function. In this case, the initial mutation must maintain enzyme activity above the threshold required for viral replication or both mutations must occur simultaneously. (B) Enabling mutations increase enzyme activity providing a buffer that permits the accumulation of mutations that on their own might lower enzyme activity below the level required for viral replication. (C) Hyperactive mutations can also directly increase tolerance to drug concentrations that would prevent the replication of WT.

The second reason that hyperactive mutations are important is that they can provide a direct growth advantage to viruses propagated in the presence of drug (Figure 1C). Based on modeling of a simple enzyme mechanism, the increase in hyperactivity of the enzyme should be directly related to the increased drug resistance for the virus. For example, a mutation that increased the catalytic rate two-fold would require twice the concentration of inhibitor to stall growth. Additionally, the impact on viral tolerance to drugs can be amplified for targets like M^pro^ that drive their own production. M^pro^ is only fully active once it cleaves itself out of its polyprotein precursors. Hyperactive mutations that increase M^pro^ activity may increase the pool of processed enzyme available for viral maturation. Consistent with this potential role of hyperactive mutations in M^pro^ resistance, selection for nirmatrelvir resistance in cell culture passaging of SARS-CoV-2 virus resulted in multiple lineages with ∼10-fold increases in tolerance to drug driven by mutations that increase enzyme activity without measurable impacts on drug binding^10,14^. However, experiments with reverse-engineered viruses harboring the most commonly observed hyperactive mutations (either T21I or L50F) showed smaller increases in IC50 in the range of 1.1-4.6 fold^14–16^.

Despite the theoretical potential of hyperactive mutants in the evolution of resistance, there has been little experimental investigation of these mutations. To further investigate the potential roles of hyperactive mutations in drug resistance, we used a mutational scanning approach to comprehensively identify mutations in M^pro^ with increased enzyme activity. We identified 175 mutations that cause significant increases in M^pro^ activity and are distributed throughout the M^pro^ structure. Viruses selected for nirmatrelvir resistance in SARS-CoV-2 passaging experiments^14–16^ were enriched for hyperactive mutations and additionally, many hyperactive mutations have also been observed in sequenced clinical SARS-CoV-2 isolates.

Therefore, hyperactive mutations appear likely to play a key role in the natural evolution of drug resistance.

## Results and Discussion

### Systematic identification of hyperactive mutations

To systematically identify all hyperactive M^pro^ mutations we used an *in vivo* fluorescent yeast reporter assay developed in our previous work^17^. For this assay, we placed the Nsp4/5 M^pro^ cut site between a CFP and YFP FRET pair such that upon expression of M^pro^ we observed a large decrease in FRET signal (Figure 2A&B). In our initial study^17^, we expressed M^pro^ to a level where WT cleavage was essentially complete in order to focus on mutations that strongly decreased enzyme activity. To identify hyperactive mutations, in this work, we decreased the duration of expression such that WT M^pro^ only partially cleaved the substrate. We monitored bulk fluorescence in reporter yeast as a function of induction time for WT M^pro^ (Figure 2B); after 40 minutes of induction we observed a clear and partial decrease in reporter signal.

**Figure 2.**
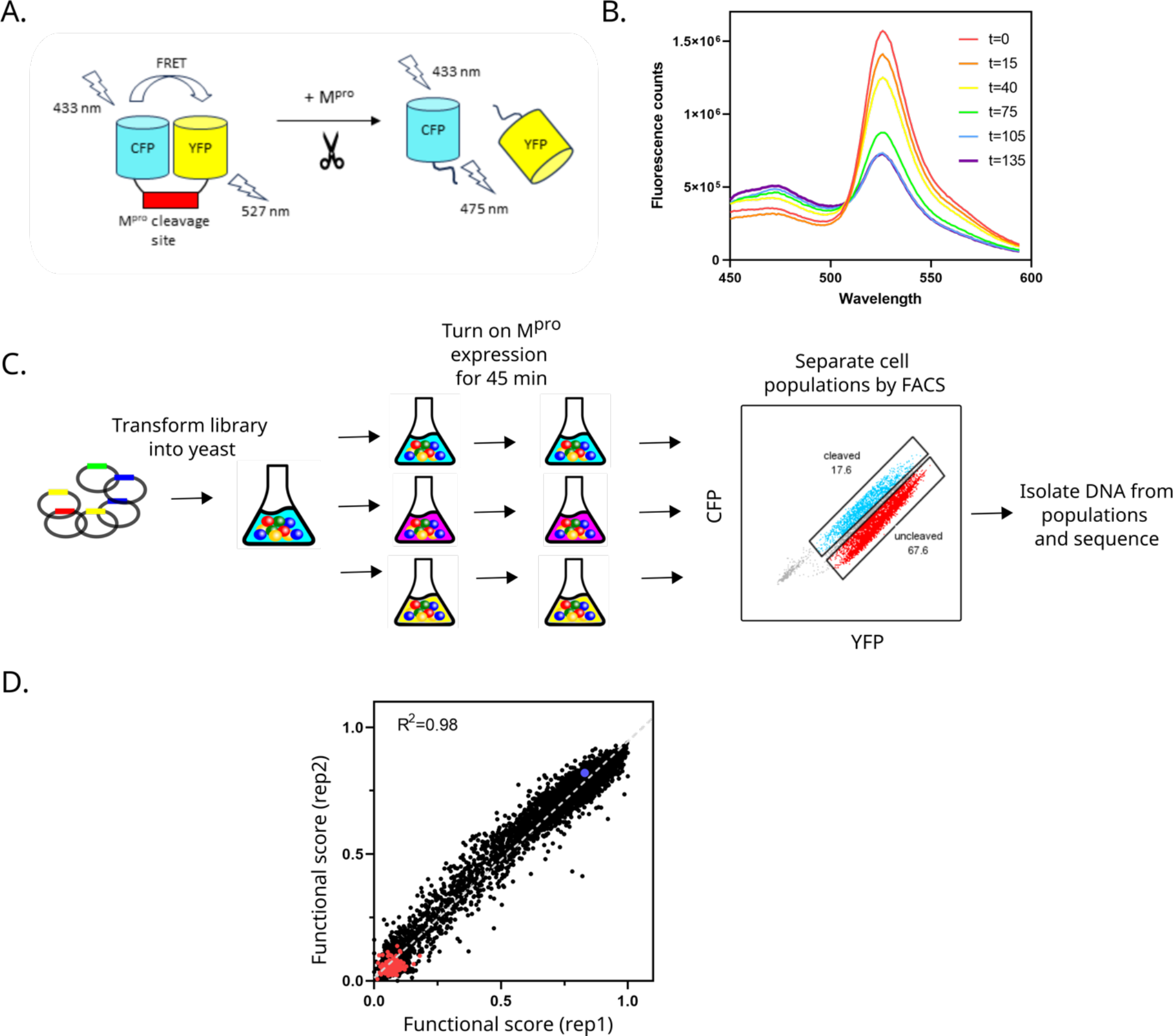
Experimental strategy to identify hyperactive M^pro^ mutants. (A) Fluorescence resonance energy transfer based reporter screen. M^pro^ variants were screened based on their ability to cleave at the Nsp4/5 cut-site, separating the YFP-CFP FRET pair. (B) Reporter cells were transformed with a plasmid expressing WT M^pro^ under the control of the LexA promoter. Cell were induced with 125 nM β-estradiol, and fluorescence assessed at the indicated timepoints. (C) Reporter cells were transformed with a plasmid library of M^pro^ mutations and split into triplicate cultures. Cells were collected after 45 minutes of induction and separated by FACS into primarily cleaved and un-cleaved populations. Each population was analyzed using a next-generation sequencing readout. (D) Correlation between functional scores of mutants from two replicates. Stop codons are shown in red and WT M^pro^ in blue.

For experiments with mutant libraries, we selected an M^pro^ induction time of 45 minutes so that we could observe partial cleavage by WT M^pro^ with the potential to identify hyperactive variants (Figure 2C). When we analyzed the resulting mutant library by flow cytometry we observed populations with varied ratios of CFP to YFP fluorescence (Figure 2D) indicating distinct levels of substrate cleavage. Because the fluorescence separation by flow cytometry was modest, we chose to sort library cells into two windows corresponding to mostly un-cleaved FRET pair (low CFP to YFP ratio) and mostly cleaved FRET pair (high CFP to YFP ratio). As in our previous work^17^, we calculated a functional score as the fraction of variant sequence reads in the cut window divided by the sum of the fraction of variant reads in the cut and uncut windows. To assess the reproducibility of our functional measurements, we performed three experimental replicates of the FACS and sequencing. Across all variants analyzed we observed a strong correlation between replicate experiments (Figure 2D and Table S1).

WT M^pro^ had a functional score of 0.83 indicating that most yeast cells harboring WT M^pro^ were in the cleaved sort window. This is consistent with partial cleavage observed in bulk fluorescence under similar conditions (Figure 2B), and small cell-to-cell variation such that the flow cytometry profile of WT moves mostly clear of the un-cleaved window. These observations suggest that additional and narrower sort windows may result in improved resolution of functional scores. However, the strong reproducibility of our functional measurements indicated that we were able to clearly identify many variants with increased activity in our assay compared to WT (Figure 2D). We used experimental replicates to identify variants that were significantly hyperactive based on a student T-test. Using this approach, we identified 175 hyperactive variants (Table S2).

### Biochemical and structural characterization of hyperactive mutants

We assessed the biochemical properties of three variants identified as hyperactive in our screen (Figure S1). Each of these variants (T21I, L50F and L141R) as well as WT M^pro^ were expressed and purified and assayed for enzyme activity using a fluorogenic Nsp4/5 peptide substrate. We chose to analyze T21I and L50F both because of their elevated functional score in our screen and because these mutations have both been selected for in SARS-CoV-2 virusultured in the presence of M^pro^ inhibitors^14^. In purified form, T21I, L50F and L141R M^pro^ all showed elevated activity (1.6, 1.7, and 1.9 fold increased enzyme proficiency compared to WT). The enzyme activity of L50F and T21I have been reported by others, and also found to be increased by a small amount (1.6 fold and 1.3 fold respectively) compared to WT^18,19^. The activity of L50F is somewhat controversial as two studies reported null activity^15,16^ and one report was unable to purify this variant^18^. In our hands, expression and purification of this variant were straightforward when cold temperatures were avoided. Corroboration of the elevated activity of T21I, L50F, and L141R in purified form demonstrates that our screen can successfully identify known hyperactive variants. Of note, T21I and L50F were both selected in multiple lineages in response to M^pro^ inhibition in culture^14–16^. T21I and L50F were at high frequency (>75%) in some lineages without any other mutations at greater than 50% levels, suggesting that they can contribute to initial resistance evolution. Both T21I and L50F also evolved to high frequency in combination with other mutations including E166V that can disrupt M^pro^ binding to inhibitors. Together these observations indicate that hyperactive mutations can both contribute to initial adaptation to M^pro^ inhibitors and can combine with other mutations to lead to higher level resistance.

We searched for hyperactivity hot-spots by counting the number of hyperactive mutations from our screen at each amino acid position in M^pro^ (Figure 3). There were about 100 positions with a single hyperactive mutation, but there were also a few positions with many hyperactive mutations. To assess the significance of these potential hot-spots, we used a boot-strap approach based on random sampling simulations. Positions with five or greater hyperactive mutations rarely occurred by random sampling and were therefore considered significant in our experimental results. There were three positions (1, 141, 151) where most amino acid changes led to hyperactivity, suggesting that the WT amino acid at these positions is poorly suited to cleavage of the substrate in our screen. In the crystal structure of catalytically-inactive M^pro^ with the Nsp4/5 substrate^20^, L141 and N151 are in local environments that are poorly suited to their biophysical properties (Figure 3B&C). The hydrophobic side-chain of L141 packs against the hydrophilic carbonyl atoms at the end of an alpha-helix. The hydrogen bonding along alpha-helices leads to a dipole where charge accumulates at the ends favoring interaction with water or other polar atoms compared to hydrophobic atoms^21,22^. In a similar biophysical mismatch, the side chain of N151 is oriented towards a core region of M^pro^ that is largely hydrophobic. In contrast to L141 and N151, S1 is not making any obviously unfavorable physical interactions in the substrate-bound structure of M^pro^. However, S1 is located in a spacious cavity where larger side-chains would have opportunities for additional favorable contacts. In addition to its role in M^pro^ activity, position 1 is the site where M^pro^ cuts itself out of viral polyprotein. Autoproteolysis is a critical step for the virus and serine at the P1’ position makes key contacts as a substrate that are important for cleavage. Consistent with its important role in substrate recognition, serine is located at the P1’ position in 6 of the 11 sites that are cleaved by M^pro^.

**Figure 3.**
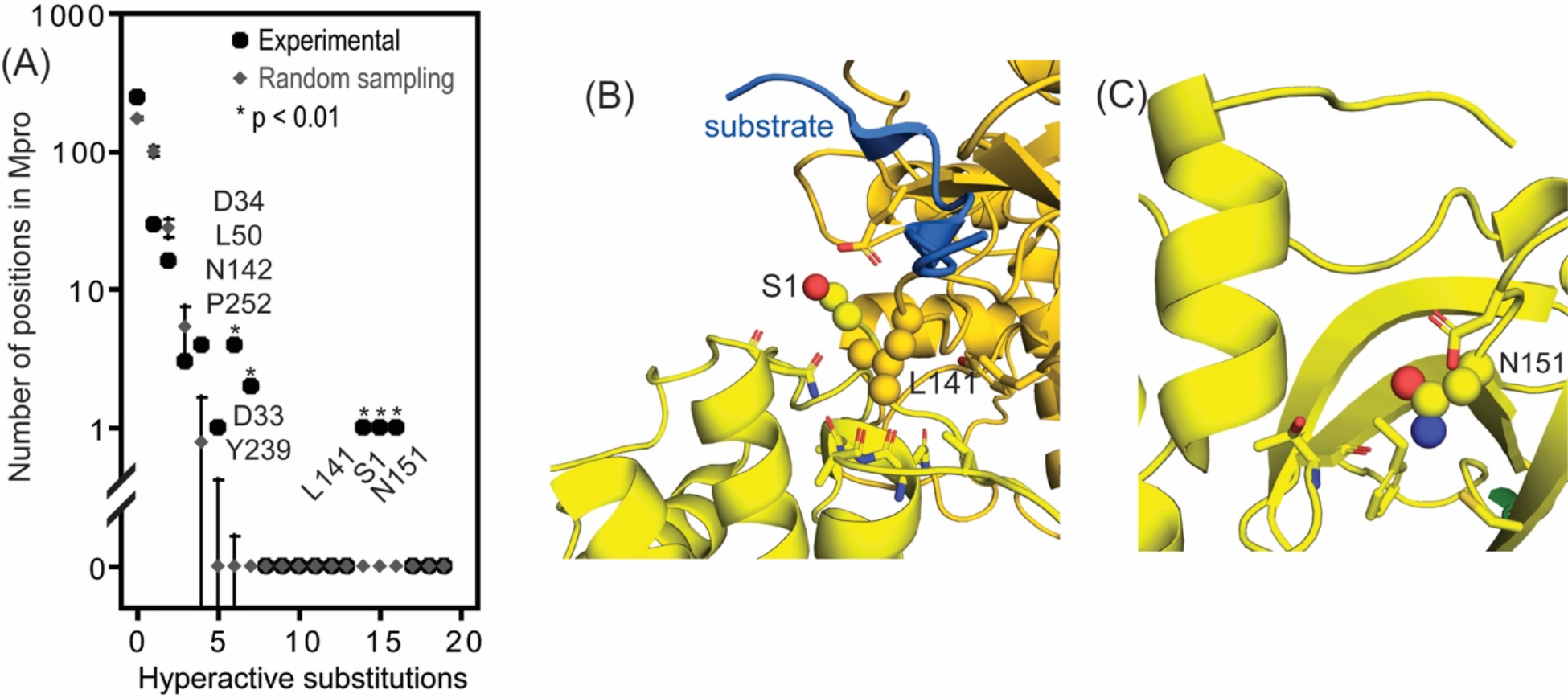
The majority of substitutions at three positions increase the functional score of M^pro^. (A) Clustering of substitutions identified as hyperactive in the FRET screen at positions in M^pro^ compared to random sampling. At three positions (1, 141, and 151), most substitutions resulted in increased functional scores. (B&C) Molecular models of M^pro^ based on the 7t70 crystal structure determined with bound substrate^20^ that illustrate the physical environment of positions 1, 141, and 151.

While selection for Ser at position 1 in M^pro^ can be explained by autoproteolysis, the poor biophysical fits of L141 and N151 in the structure are likely related to their roles in enzyme activity. Hyperactive mutations at L141 and N151 are present at low frequency (<0.1%) in sequenced isolates of SARS-CoV-2 (Figure S2). Of note, phylogenetic analyses of sequenced isolates suggest that mutations at positions 141 and 151 of M^pro^ cause fitness defects^23^. We considered the possibility that hyperactive mutations are deleterious when drug is not present (as has been observed for L50F^16^), but did not observe any meaningful differences in the frequency in sequenced isolates of mutations with WT-like from those with increased enzyme activity. Of note, the relationship between enzyme activity and fitness are frequently non-linear^24–26^ and can be complex, particularly for viral enzymes including proteases that act on more than one substrate^27^. Additionally, M^pro^ expression is known to be toxic in many host cells, suggesting the potential for selection beyond viral proteins^28,29^. Future studies of the impacts of M^pro^ mutations on additional cut-sites may help to clarify the relationship between enzyme activity and viral fitness.

We also identified a handful of hot-spot positions where 6 or 7 mutations were hyperactive in our yeast screen (D33, D34, L50, N142, Y239 and P252) (Figure 3A). Two of these positions (50 and 252) have been observed in multiple lineages selected for nirmatrelvir resistance in cell culture^14^. All of the hyperactive mutations at 252 (P to C, F, L, M, V, or Y) were hydrophobic, suggesting that this is a key feature mediating activity at this position. For the other hot-spots, there did not appear to be a single biophysical feature shared by all the hyperactive variants. For example, at L50, mutations to aromatic amino acids (F, H, Y) or a subset of polar amino acids (Q, S, T) were hyperactive in our yeast screen.

To explore the mechanism of hyperactive mutations, we determined the structure of three variants at hyperactive hot-spots (L50F, L141R, and N142P). We chose to investigate L50F because it was observed in multiple nirmatrelvir-selected lineages in cell culture^14^. L141R and N142P were selected because they are both close to the active site and are dramatic amino acid changes which may lead to clear structural changes. We were able to readily grow diffraction-quality crystals of all three variants in complex with inhibitors (nirmatrelvir for L50F and PF-00835231 for L141R and N142P). The structures of each variant were aligned closely to WT with some noticeable changes around the active site (Figure 4). The most notable change in the structure of L50F from the WT structure was a ∼1 angstrom shift of backbone atoms in residues 1-6 and 290-298 of the other monomer, causing the amide of M6 to form a closer H-bond with the carbonyl oxygen of R4. Of note, R4 and M6 are more than 20 angstroms from the L50F mutation, indicating that physical changes can be transmitted widely across the M^pro^ structure. Differences between WT and L141R were more concentrated local to the mutation. The mutated arginine at position 141 forms multiple hydrogen bonds with shifts in the position of residues that form the new hydrogen bonds. The main chain of N142P aligns very closely with WT suggesting that this mutation does not dramatically alter the WT inhibitor-bound to inhibitor. Altogether, these analyses suggest that subtle structural changes can lead to the modest hyperactivity phenotypes we have identified.

**Figure 4.**
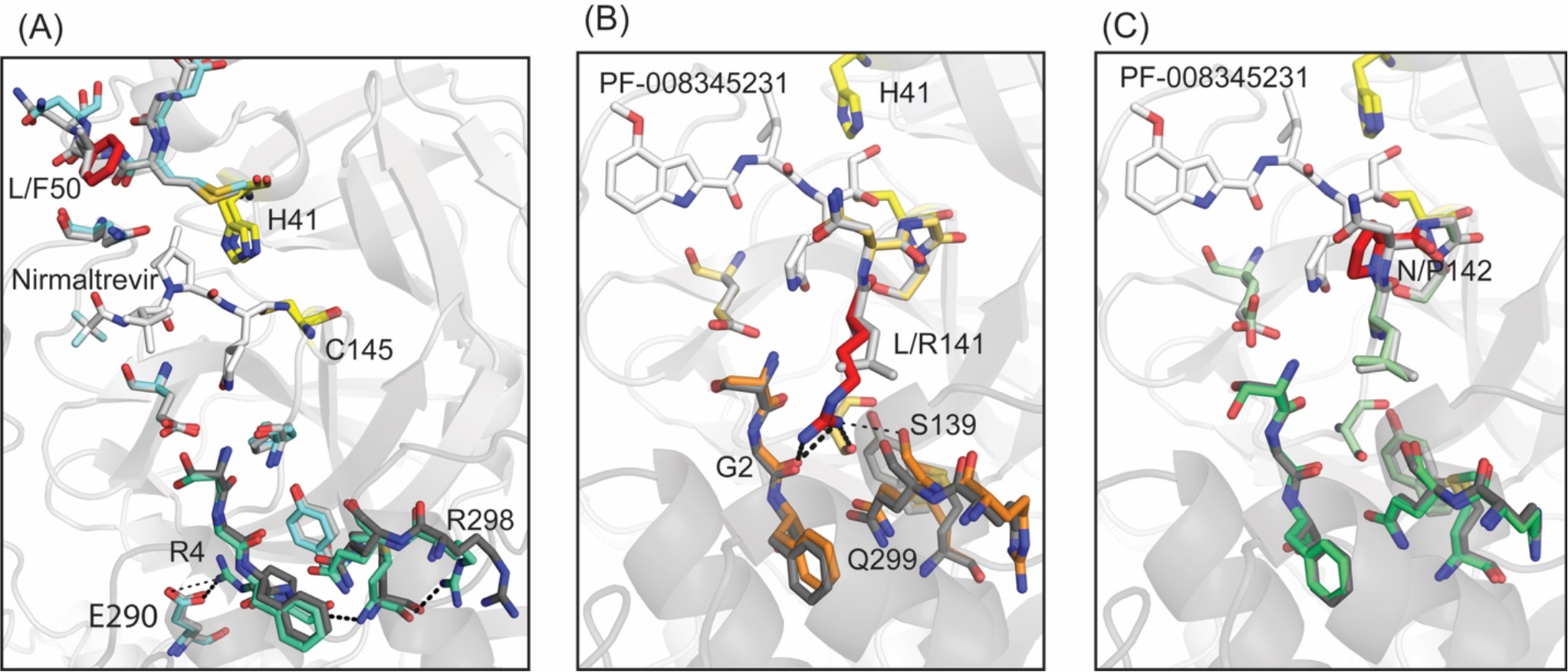
Comparison of the experimentally determined X-ray crystal structures of individual hyperactive variants to WT M^pro^. The mutated side-chains are colored red in all panels. (A) Overlay of WT M^pro^ (7RFS.PDB) with the structure of L50F. WT side chains are shown in grey or black and L50F side chains in cyan, pale green, or red. (B&C) WT M^pro^ overlayed with either L141R (B) or N142P (C). WT side chains are shown in grey or black and mutant side chains in red and orange (B), or red and green (C).

We examined the physical distribution of all of the hyperactive mutations that we identified in our screen (Figure 5). Of note, most of these mutations did not occur at hot-spots (positions with five or more hyperactive mutations). Instead, the hyperactive mutations we identified were located broadly throughout the structure of M^pro^ (Figure 5A). Given this broad distribution, we searched for properties of positions in the structure where hyperactive mutations might occur preferentially (Figure 5B). We used a bootstrap approach to randomly sample positions in M^pro^ to estimate expected variation and significance. Almost all properties showed hyperactive mutations at similar frequencies compared to all positions in M^pro^, indicating that these properties (solvent accessibility, distance to substrate, location at interfaces) do not correlate with or predict hyperactivity. The only property that was significantly different than random expectations was location in a helix where hyperactive mutations were less prevalent than random. The inability to accurately predict hyperactivity from fixed structures may indicate that protein dynamics may be a primary driver for hyperactivity. The impacts of protein motions on enzyme function are notoriously difficult to analyze^30^ and remain one of the most important and understudied areas of biochemistry. However, where they have been analyzed, protein dynamics have been shown to have critical impacts on enzyme activity. Interestingly, the one feature of M^pro^ that we found significantly depleted for hyperactive mutations was location in a helix. Of all secondary structure elements, helices are known to be the least perturbed by mutation^31^. The contribution of dynamics to hyperactive mutations in M^pro^ will require future efforts to resolve with clarity.

**Figure 5.**
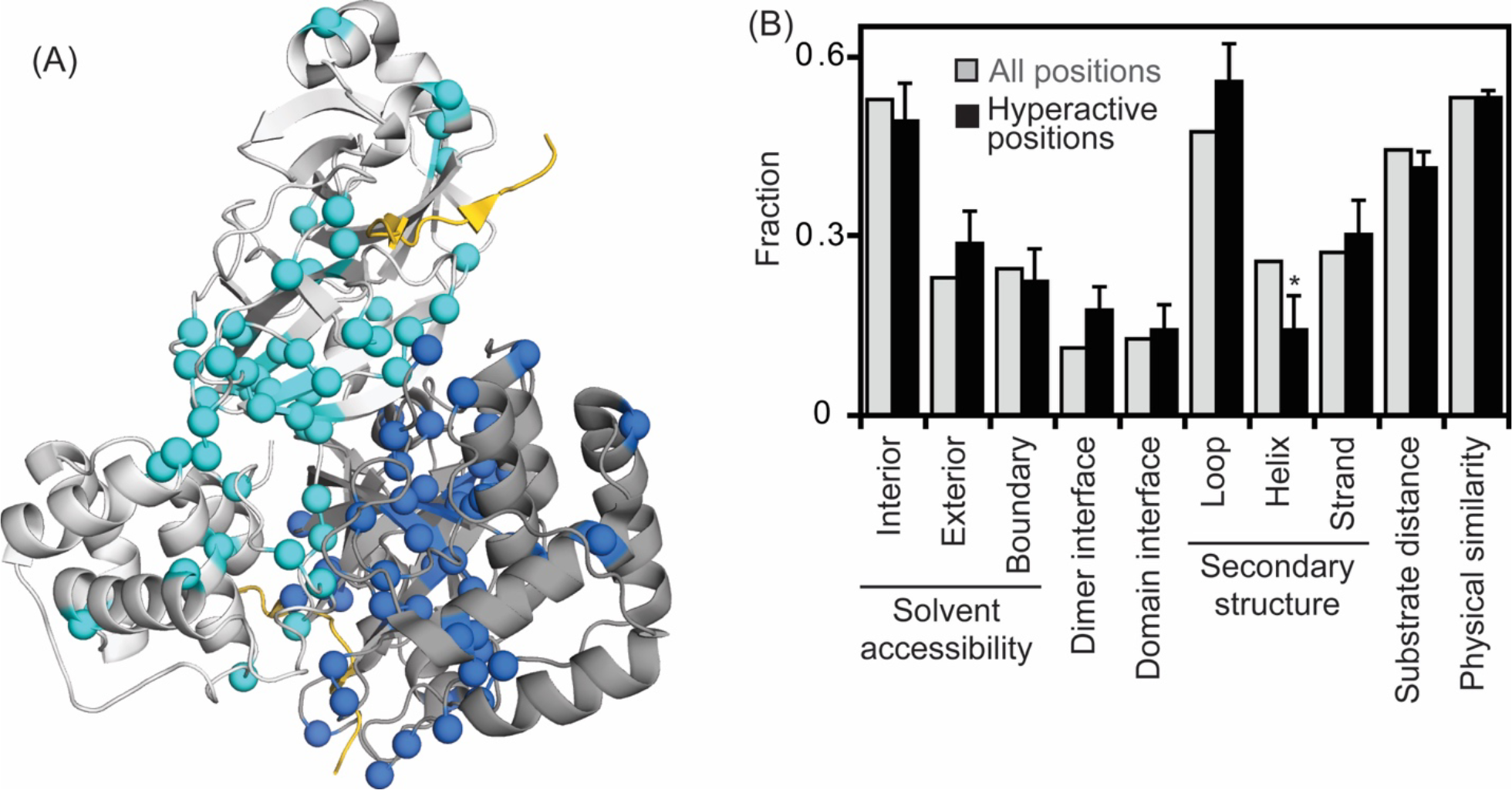
Structural and physical characteristics of positions with hyperactive mutations compared to all positions in M^pro^. (A) Structural representation of Mpro based on the 7t70.PDB^20^ crystal structure showing the location of all identified hyperactive mutations (colored in blue) and bound substrate in yellow. Shading was used to distinguish each subunit in the dimer. (B) Properties of positions with hyperactive mutations. The only property that was statistically significant (p<0.05 indicated with a *) for hyperactive positions was location in a helix. Error bars were estimated by bootstrapping and significance was calculated from Z-scores.

### Occurrence of hyperactive mutants in natural isolates and cell culture-selected lineages

Motivated by the potential role of both hyperactive mutations and drug binding mutations in the evolution of drug resistance in SARS-CoV-2, we surveyed their prevalence (Figure 6). In Figure 6A, we examined the prevalence in viruses selected for nirmatrelvir resistance in culture^14–16^. Most mutations that evolved in cell culture showed either hyperactivity or reduced drug binding, consistent with these two features as the main drivers of resistance. Interestingly, mutations that we identified as hyperactive were about three times as prevalent in drug selected lineages compared to mutations we previously identified as drug binding in similar mutational scans^10^. The fraction of all point variants that were either hyperactive or reduced drug binding was far smaller than the mutations in these categories in drug-selected viral lineages (Figure 6B). These findings strongly indicate that selection rather than random chance caused enrichment of these categories of mutations in the drug-selected lineages.

**Figure 6.**
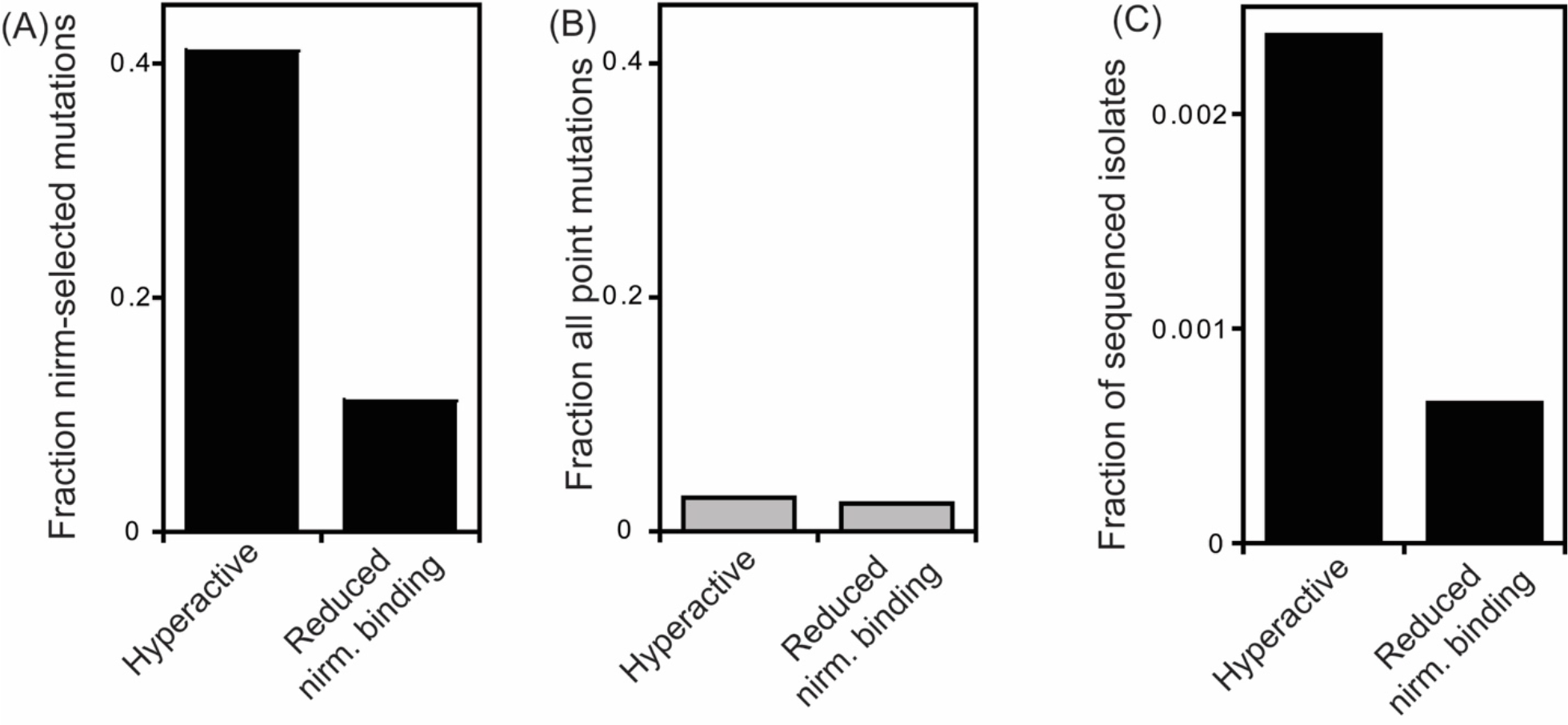
Hyperactive mutations are more frequently observed than drug binding mutations in viruses selected for nirmatrelvir resistance and in sequenced isolates. (A) Graph of the fraction of substitutions identified in viruses selected for nirmatrelvir resistance^14^ that exhibited M^pro^ hyperactivity (this work), or reduced nirmatrelvir binding^10^. (B) Graph illustrating the relative fraction of all possible mutations that were identified as either hyperactive or having reduced nirmatrelvir binding. (C) The fraction of sequenced SARS-CoV-2 isolates^32^ with mutations in M^pro^ identified as either hyperactive or with reduced nirmatrelvir binding.

To explore the potential mechanistic roles of hyperactive mutations in response to drug selection (Figure 1), we examined their occurrence with mutations identified as reducing drug binding^10^ in cell culture-selected lineages^14^. In the drug-selection experiments, most mutations that were reported had arisen close to fixation (110 mutations at a frequency >75%, 7 at 50-75%, 15 at 25-50%, and 11 at 5-25%), indicating a prevalence of selective sweeps of individual variants and relatively low levels of clonal interference. For this reason, we focused our analyses on mutations reported to have become the major species with frequencies over 50% in a lineage. Out of 53 selected lineages, there were 7 that only accumulated a single hyperactive mutation, 0 that had only accumulated a single reduced drug-binding mutation, and 25 that had accumulated a hyperactive and binding mutation. The observation of multiple drug-selected lineages only harboring hyperactive mutations indicates that they can increase resistance on their own. The observation of many drug-selected lineages with both hyperactive and reduced drug-binding mutations indicates that these categories of mutation can synergize to generate viruses that are fit and strongly drug resistant.

Next, we examined the prevalence of hyperactive and drug-binding mutations in sequenced isolates. While mutations that we identified in these categories were observed in circulating SARS-CoV-2 variants, they were at very low levels (Figure 6C). In addition, they remain at low levels in viruses that have been sequenced in recent months (Figure S3). These findings indicate that the clinical use of M^pro^ inhibitors has not yet resulted in a high frequency of mutations that can contribute to drug resistance. The similar level of hyperactive and drug-binding mutations in recent viral isolates compared to isolates from throughout the pandemic suggests that they predominantly represent standing variation – mutations whose frequency is not yet driven by positive selection. The low prevalence of mutations that can contribute to resistance indicates that the efficacy of M^pro^ inhibitors is largely uncompromised by resistance mutations. However, the low-level presence of these mutations also means that the viral population could readily respond to widespread use of M^pro^ inhibitors by increasing the frequency of resistant mutations. Interestingly, hyperactive mutations are roughly three times as prevalent in sequenced isolates as reduced drug-binding mutations. This may indicate that hyperactive mutations have a fitness advantage compared to mutations that reduce drug binding, consistent with the tendency of drug-binding mutations to also reduce enzyme activity^13^.

## Conclusions

We have systematically analyzed hyperactive mutations in M^pro^. Our results indicate that hyperactive mutations are roughly as common as mutations that reduce drug binding in M^pro^. However, hyperactive mutations are disproportionately observed in cell culture lineages selected for resistance and in the standing variation of sequenced isolates, indicating that hyperactive mutations are likely to play a key role in the evolution of resistance in circulating variants. The observation of hyperactive mutations as the only high frequency change in some resistant lineages selected in cell culture indicates that they can directly contribute to resistance, consistent with biochemical principles (Figure 1). Hyperactive mutations can also contribute to resistance by either enabling or rescuing mutations that disrupt drug binding that commonly also reduce enzyme activity (Figure 1). Our systematic analysis of hyperactive mutations enables future efforts to efficiently survey and detect early stages of drug resistance in circulating variants.

There are many important issues that this work raises that will be the focus of future efforts. These include generating a publicly available surveillance record of mutations that can cause resistance in sequenced SARS-CoV-2 isolates, and detailed evolutionary models to investigate how different levels of M^pro^ inhibitors can impact resistance evolution.

## Materials and Methods

### Generation of Mutant Libraries

The SARS-CoV-2 single site library was synthesized by Twist Biosciences (www.twist.com) with each amino acid position modified to all 19 amino acid positions plus a stop codon. The libraries were cloned and barcoded as described previously^17^. In short, the library was fused to a Ubiquitin gene and assembled into a destination vector. The Ubiquitin-M^pro^ fusion protein is cleaved *in vivo* by Ub-specific proteases directly C-terminal to the Ub, allowing expression of M^pro^ with its authentic N-terminal serine residue^33^. To precisely control the expression level of M^pro^ while limiting its toxic side effects, the Ub-M^pro^ library was placed under control of the β-estradiol-activated LexA-ER-AD transcription factor^34^. The cloned library was barcoded with a randomized 18 bp nucleotide sequence and the barcodes were associated with the M^pro^ variants using PacBio sequencing. The barcoded plasmid mutant library was combined with a plasmid containing wild-type M^pro^ associated with approximately 150 unique barcodes.

### Bulk FRET competition experiment

The FRET reporter strain was constructed by fusing together the two fluorescent proteins, YPet and CyPet, with the Nsp4/5 M^pro^ cleavage site engineered in the middle with a linker of two glycines as previously described (YPet-GGTSAVLQ/SGFRKMGG-CyPet). The YPet-CyPet FRET pair is a YFP-CFP fluorescent pair that has been fluorescently optimized by directed evolution for intracellular FRET^35^. The resulting YPet-nsp4/5-CyPet fusion gene was integrated into the *ade2* gene locus of W303 cells (*leu2-3,112 trp1-1 can1-100 ura3-1 ade2-1 his3-11,15*). The blended WT and mutant plasmid library was transformed using the LiAc/PEG procedure^36^ into the FRET reporter strain. To avoid bottlenecking, sufficient transformation reactions were performed to achieve approximately 5 million independent yeast transformations representing a 50-fold sampling of the average barcode. Following 12 hours of recovery in synthetic dextrose media lacking adenine (SD-A), transformed cells were washed three times in SD lacking adenine and uracil (SD-A-U) (to select for the presence of the M^pro^ variant plasmid) and grown in 500 mL SD-A-U media at 30°C for 48 hr with repeated dilutions to maintain the cells in log phase growth and expand the library. To avoid population bottlenecks, at least 10^7^ cells were passed for each dilution. Next, the library was diluted to early log phase in SD-A-U, grown for 3 hours, and 125 nM β-estradiol (from a 10 mM stock in 95% ethanol, Sigma-Aldrich) was added to induce M^pro^ expression. Cultures were grown with shaking at 180 rpm for 45 min, at which time samples of ∼10^7^ cells were collected. The cells were immediately washed three times with ice cold tris-buffered saline containing 0.1% Tween and 0.1% bovine serum albumin (TBST-BSA), diluted to 10^6^ cells/mL and immediately sorted for YFP and CFP expression on a FACS Aria II cell sorter. Cells expressing the cut FRET pair (low FRET) were collected in one population and the uncut FRET pair (high FRET) in a second population. We collected 5 million cells of each population to ensure adequate library coverage. Yeast cells were collected by centrifugation and stored at - 80°C. We performed three technical replicates for this experiment in which the library was transformed into yeast cells in bulk and then separated into three flasks. For each replicate, cell growth, FACS separation, and sequence analyses were performed independently.

### DNA preparation and sequencing

We isolated plasmid DNA from each FACS cell population for each replicate as described^24^. Purified plasmid DNA was linearized with AscI (NEB). Barcodes were amplified with 25 cycles of PCR using the entire sample of extracted DNA from each cell population to ensure complete sequencing coverage. PCR was performed using Phusion polymerase (NEB) and primers that add Illumina adapter sequences and a 6 bp identifier sequence used to distinguish cell populations. PCR products were purified two times over silica columns (Zymo Research) and the DNA concentration was measured by qPCR using the KAPA SYBR FAST qPCR Master Mix (Kapa Biosystems) on a Bio-Rad CFX machine. PCR samples were combined and sequenced on an Illumina NextSeq instrument using a NextSeq 500/550 High Output Kit v. 2.5 (75 cycles). The Illumina barcode reads were calculated using custom scripts that have been deposited on GitHub (https://github.com/JuliaFlynn/BolonLab). Reads were filtered for Phred scores > 10 and strict matching to the expected template and identifier sequences. Filtered reads were parsed based on their identifier sequences. For each identifier sequence, each unique N18 barcode read was counted. Using the variant-barcode association table that was generated by PacBio sequencing^17^, the frequency of each mutant in each cell population was tabulated. To calculate the functional score for each mutant, the fraction of each mutant in the cut and uncut windows was first calculated by dividing the sequencing count of each mutant in a window by the total counts in that window. The functional score was then calculated as the fraction of the mutant in the cut window divided by the sum of the fraction of the mutant in the cut and uncut windows.

### Purification of M^pro^

His_6_-SUMO-SARS-CoV-2 M^pro^ (WT) was cloned into a pETite vector^20^. L50F, L141R and N142P mutations were created by site-specific mutagenesis. The vectors were transformed into Hi-Control BL21 (DE3) *Escherichia coli cells* and the M^pro^ proteins were expressed and purified as previously described^20^.

### Analyses of enzyme properties of purified M^pro^

To measure the K_m_ of M^pro^ mutants, 0.1 μM enzyme was added to a series of 0-125 μM FRET substrate (Dabcyl-KTSAVLQSGFRKME-Edans (LifeTein)) in 50 mM Tris pH 7.5, 50 mM NaCl, 1 mM EDTA, 1 mM DTT and 4% DMSO. The cleavage reaction was monitored using a PerkinElmer EnVision plate reader at 30°C (355 nM excitation, 460 nM emission). Three replicates were performed with each mutant. The absorbance of the FRET substrate at the highest concentration (125 μM) was less than 0.1, and so the interference due to the inner filter effect was assumed to be minimal. The initial velocity was plotted against FRET substrate concentrations and were fit using GraphPad Prism 9 to the Michaelis-Menten equation.

### Co-crystallization

Pfizer compounds PF-07321332 and PF-00835231 were purchased from Selleckchem.com. Protein complexes were made by incubating 6-7 mg/mL of each mutant M^pro^ with 5-10 molar excess small molecule in 20 mM HEPES pH 7.5, 300 mM NaCl buffer on the benchtop for one hour. Co-crystals were obtained with 10-20 % (w/v) PEG 3350, 0.20-0.30 M NaCl, and 0.1 M Bis-Tris-Methane pH 5.5 using pre-greased, 24-well VDX hanging drop trays (Hampton Research Corporation). Diffraction quality crystals required 10 days, on average, of growth time at room temperature. Varying the protein complex to crystallization solution ratios (1 μL:2 μL, 2 μL:2 μL, and 3 μL:2 μL) helped obtain large, singular crystals. Microseeding was not required for crystal growth.

### Data Collection and Structure Determination

Prior to data collection, crystals were soaked in their respective mother liquor solutions supplemented with 25 % glycerol. Diffraction data was collected under cryogenic conditions (100 K) at the University of Massachusetts Chan Medical School X-Ray Crystallography and Drug Design Core facility (Worcester, MA). Home source data acquisition was performed using a Rigaku MicroMax-007HF x-ray generator equip with a HyPix-6000HE detector. Data processing (indexing, integration, and scaling) was done using CrysAlis^Pro^PX (Rigaku Corporation). Structure solutions, model building, and refinement were done using the Phenix suite of software packages^37^. Molecular replacement solutions were obtained with PHASER^38^ using PDB 7L0D as the reference model. Model building and structure refinement were performed using COOT^39^. Reference model bias was minimized by reserving 5% of the collected data for R_free_ calculations^40^. Data collection parameters and refinement statistics are available in Table S3.

### Identifying mutations in circulating SARS-CoV-2 sequences

The frequency of M^pro^ variants was investigated based on high-quality SARS-CoV-2 sequence data^32^. We downloaded the latest variable call format file (public-latest.all.masked.vcf.gz) from http://hgdownload.soe.ucsc.edu/goldenPath/wuhCor1/UShER_SARS-CoV-2/ on Sept 7, 2023. We wrote a custom perl script to tabulate sequence observations of mutations identified as hyperactive in this work, or with reduced binding to nirmaltrevir or ensitrelvir based on a previously published yeast screen^10^. The frequency of these classes of mutations was calculated by dividing by the total number of sequenced isolates in the time window analyzed.

## References

(1) Dadgostar, P. Antimicrobial Resistance: Implications and Costs. Infect. Drug Resist. 2019, Volume 12, 3903–3910. 10.2147/IDR.S234610.

(2) Chinemerem Nwobodo, D.; Ugwu, M. C.; Oliseloke Anie, C.; Al-Ouqaili, M. T. S.; Chinedu Ikem, J.; Victor Chigozie, U.; Saki, M. Antibiotic Resistance: The Challenges and Some Emerging Strategies for Tackling a Global Menace. J. Clin. Lab. Anal. 2022, 36 (9), e24655. 10.1002/jcla.24655.

(3) Strathdee, S. A.; Davies, S. C.; Marcelin, J. R. Confronting Antimicrobial Resistance beyond the COVID-19 Pandemic and the 2020 US Election. Lancet Lond. Engl. 2020, 396 (10257), 1050–1053. 10.1016/S0140-6736(20)32063-8.

(4) Herold, J.; Raabe, T.; Schelle-Prinz, B.; Siddell, S. G. Nucleotide Sequence of the Human Coronavirus 229E RNA Polymerase Locus. Virology 1993, 195 (2), 680–691. 10.1006/viro.1993.1419.

(5) Lim, K. P.; Ng, L. F.; Liu, D. X. Identification of a Novel Cleavage Activity of the First Papain-like Proteinase Domain Encoded by Open Reading Frame 1a of the Coronavirus Avian Infectious Bronchitis Virus and Characterization of the Cleavage Products. J. Virol. 2000, 74 (4), 1674–1685. 10.1128/jvi.74.4.1674-1685.2000.

(6) Ziebuhr, J.; Herold, J.; Siddell, S. G. Characterization of a Human Coronavirus (Strain 229E) 3C-like Proteinase Activity. J. Virol. 1995, 69 (7), 4331–4338. 10.1128/JVI.69.7.4331-4338.1995.

(7) Jin, Z.; Du, X.; Xu, Y.; Deng, Y.; Liu, M.; Zhao, Y.; Zhang, B.; Li, X.; Zhang, L.; Peng, C.; Duan, Y.; Yu, J.; Wang, L.; Yang, K.; Liu, F.; Jiang, R.; Yang, X.; You, T.; Liu, X.; Yang, X.; Bai, F.; Liu, H.; Liu, X.; Guddat, L. W.; Xu, W.; Xiao, G.; Qin, C.; Shi, Z.; Jiang, H.; Rao, Z.; Yang, H. Structure of Mpro from SARS-CoV-2 and Discovery of Its Inhibitors. Nature 2020, 582 (7811), 289–293. 10.1038/s41586-020-2223-y.

(8) Rufer, A. C. Drug Discovery for Enzymes. Drug Discov. Today 2021, 26 (4), 875–886. 10.1016/j.drudis.2021.01.006.

(9) Nalam, M. N.; Schiffer, C. A. New Approaches to HIV Protease Inhibitor Drug Design II: Testing the Substrate Envelope Hypothesis to Avoid Drug Resistance and Discover Robust Inhibitors: Curr. Opin. HIV AIDS 2008, 3 (6), 642–646. 10.1097/COH.0b013e3283136cee.

(10) Flynn, J. M.; Huang, Q. Y. J.; Zvornicanin, S. N.; Schneider-Nachum, G.; Shaqra, A. M.; Yilmaz, N. K.; Moquin, S. A.; Dovala, D.; Schiffer, C. A.; Bolon, D. N. A. Systematic Analyses of the Resistance Potential of Drugs Targeting SARS-CoV-2 Main Protease; preprint; Molecular Biology, 2023. 10.1101/2023.03.02.530652.

(11) Gulnik, S. V.; Suvorov, L. I.; Liu, B.; Yu, B.; Anderson, B.; Mitsuya, H.; Erickson, J. W. Kinetic Characterization and Cross-Resistance Patterns Of HIV-1 Protease Mutants Selected under Drug Pressure. Biochemistry 1995, 34 (29), 9282–9287. 10.1021/bi00029a002.

(12) Lin, Y.; Lin, X.; Hong, L.; Foundling, S.; Heinrikson, R. L.; Thaisrivongs, S.; Leelamanit, W.; Raterman, D.; Shah, M. Effect of Point Mutations on the Kinetics and the Inhibition of Human Immunodeficiency Virus Type 1 Protease: Relationship to Drug Resistance. Biochemistry 1995, 34 (4), 1143–1152. 10.1021/bi00004a007.

(13) Kurt Yilmaz, N.; Swanstrom, R.; Schiffer, C. A. Improving Viral Protease Inhibitors to Counter Drug Resistance. Trends Microbiol. 2016, 24 (7), 547–557. 10.1016/j.Am.2016.03.010.

(14) Iketani, S.; Mohri, H.; Culbertson, B.; Hong, S. J.; Duan, Y.; Luck, M. I.; Annavajhala, M. K.; Guo, Y.; Sheng, Z.; Uhlemann, A.-C.; Goff, S. P.; Sabo, Y.; Yang, H.; Chavez, A.; Ho, D. D. Multiple Pathways for SARS-CoV-2 Resistance to Nirmatrelvir. Nature 2023, 613 (7944), 558–564. 10.1038/s41586-022-05514-2.

(15) Jochmans, D.; Liu, C.; Donckers, K.; Stoycheva, A.; Boland, S.; Stevens, S. K.; De Vita, C.; Vanmechelen, B.; Maes, P.; Trüeb, B.; Ebert, N.; Thiel, V.; De Jonghe, S.; Vangeel, L.; Bardiot, D.; Jekle, A.; Blatt, L. M.; Beigelman, L.; Symons, J. A.; Raboisson, P.; Chaltin, P.; Marchand, A.; Neyts, J.; Deval, J.; Vandyck, K. The Substitutions L50F, E166A, and L167F in SARS-CoV-2 3CLpro Are Selected by a Protease Inhibitor In Vitro and Confer Resistance To Nirmatrelvir. mBio 2023, 14 (1), e02815–22. 10.1128/mbio.02815-22.

(16) Zhou, Y.; Gammeltoh, K. A.; Ryberg, L. A.; Pham, L. V.; Tjørnelund, H. D.; Binderup, A.; Duarte Hernandez, C. R.; Fernandez-Antunez, C.; Offersgaard, A.; Fahnøe, U.; Peters, G. H. J.; Ramirez, S.; Bukh, J.; Gottwein, J. M. Nirmatrelvir-Resistant SARS-CoV-2 Variants with High Fitness in an Infectious Cell Culture System. Sci. Adv. 2022, 8 (51), eadd7197. 10.1126/sciadv.add7197.

(17) Flynn, J. M.; Samant, N.; Schneider-Nachum, G.; Barkan, D. T.; Yilmaz, N. K.; Schiffer, C. A.; Moquin, S. A.; Dovala, D.; Bolon, D. N. Comprehensive Fitness Landscape of SARS-CoV-2 Mpro Reveals Insights into Viral Resistance Mechanisms. eLife 2022, 11, e77433. 10.7554/eLife.77433.

(18) Duan, Y.; Zhou, H.; Liu, X.; Iketani, S.; Lin, M.; Zhang, X.; Bian, Q.; Wang, H.; Sun, H.; Hong, S. J.; Culbertson, B.; Mohri, H.; Luck, M. I.; Zhu, Y.; Liu, X.; Lu, Y.; Yang, X.; Yang, K.; Sabo, Y.; Chavez, A.; Goff, S. P.; Rao, Z.; Ho, D. D.; Yang, H. Molecular Mechanisms of SARS-CoV-2 Resistance to Nirmatrelvir. Nature 2023. 10.1038/s41586-023-06609-0.

(19) Chen, S. A.; Arutyunova, E.; Lu, J.; Khan, M. B.; Rut, W.; Zmudzinski, M.; Shahbaz, S.; Iyyathurai, J.; Moussa, E. W.; Turner, Z.; Bai, B.; Lamer, T.; Nieman, J. A.; Vederas, J. C.; Julien, O.; Drag, M.; Elahi, S.; Young, H. S.; Lemieux, M. J. SARS-CoV-2 Mpro Protease Variants of Concern Display Altered Viral Substrate and Cell Host Target Galectin-8 Processing but Retain Sensitivity toward Antivirals. ACS Cent. Sci. 2023, 9 (4), 696–708. 10.1021/acscentsci.3c00054.

(20) Shaqra, A. M.; Zvornicanin, S. N.; Huang, Q. Y. J.; Lockbaum, G. J.; Knapp, M.; Tandeske, L.; Bakan, D. T.; Flynn, J.; Bolon, D. N. A.; Moquin, S.; Dovala, D.; Kurt Yilmaz, N.; Schiffer, C. A. Defining the Substrate Envelope of SARS-CoV-2 Main Protease to Predict and Avoid Drug Resistance. Nat. Commun. 2022, 13 (1), 3556. 10.1038/s41467-022-31210-w.

(21) Aurora, R.; Rosee, G. D. Helix Capping: Helix Capping. Protein Sci. 1998, 7 (1), 21–38. 10.1002/pro.5560070103.

(22) Serrano, L.; Fersht, A. R. Capping and α-Helix Stability. Nature 1989, 342 (6247), 296–299. 10.1038/342296a0.

(23) Bloom, J. D.; Neher, R. A. Fitness Effects of Mutations to SARS-CoV-2 Proteins; preprint; Evolutionary Biology, 2023. 10.1101/2023.01.30.526314.

(24) Jiang, L.; Mishra, P.; Hietpas, R. T.; Zeldovich, K. B.; Bolon, D. N. A. Latent Effects of Hsp90 Mutants Revealed at Reduced Expression Levels. PLoS Genet. 2013, 9 (6), e1003600. 10.1371/journal.pgen.1003600.

(25) Kacser, H.; Burns, J. A. THE MOLECULAR BASIS OF DOMINANCE. Genetics 1981, 97 (3–4), 639–666. 10.1093/genetics/97.3-4.639.

(26) Cisneros, A. F.; Gagnon-Arsenault, I.; Dubé, A. K.; Després, P. C.; Kumar, P.; Lafontaine, K.; Pelletier, J. N.; Landry, C. R. Epistasis between Promoter Activity and Coding Mutations Shapes Gene Evolvability. Sci. Adv. 2023, 9 (5), eadd9109. 10.1126/sciadv.add9109.

(27) Schneider-Nachum, G.; Flynn, J.; Mavor, D.; Schiffer, C. A.; Bolon, D. N. A. Analyses of HIV Proteases Variants at the Threshold of Viability Reveals Relationships between Processing Efficiency and Fitness. Virus Evol. 2021, 7 (2), veab103. 10.1093/ve/veab103.

(28) Resnick, S. J.; Iketani, S.; Hong, S. J.; Zask, A.; Liu, H.; Kim, S.; Melore, S.; Lin, F.-Y.; Nair, M. S.; Huang, Y.; Lee, S.; Tay, N. E. S.; Rovis, T.; Yang, H. W.; Xing, L.; Stockwell, B. R.; Ho, D. D.; Chavez, A. Inhibitors of Coronavirus 3CL Proteases Protect Cells from Protease-Mediated Cytotoxicity. J. Virol. 2021, 95 (14), e0237420. 10.1128/JVI.02374-20.

(29) Cao, W.; Cho, C.-C. D.; Geng, Z. Z.; Shaabani, N.; Ma, X. R.; Vatansever, E. C.; Alugubelli, Y. R.; Ma, Y.; Chaki, S. P.; Ellenburg, W. H.; Yang, K. S.; Qiao, Y.; Allen, R.; Neuman, B. W.; Ji, H.; Xu, S.; Liu, W. R. Evaluation of SARS-CoV-2 Main Protease Inhibitors Using a Novel Cell-Based Assay. ACS Cent. Sci. 2022, 8 (2), 192–204. 10.1021/acscentsci.1c00910.

(30) SAller, J. B.; Otten, R.; Häussinger, D.; Rieder, P. S.; Theobald, D. L.; Kern, D. Structure Determination of High-Energy States in a Dynamic Protein Ensemble. Nature 2022, 603 (7901), 528–535. 10.1038/s41586-022-04468-9.

(31) Abrusán, G.; Marsh, J. A. Alpha Helices Are More Robust to Mutations than Beta Strands. PLOS Comput. Biol. 2016, 12 (12), e1005242. 10.1371/journal.pcbi.1005242.

(32) McBroome, J.; Thornlow, B.; Hinrichs, A. S.; Kramer, A.; De Maio, N.; Goldman, N.; Haussler, D.; Corbett-Detig, R.; Turakhia, Y. A Daily-Updated Database and Tools for Comprehensive SARS-CoV-2 Mutation-Annotated Trees. Mol. Biol. Evol. 2021, 38 (12), 5819–5824. 10.1093/molbev/msab264.

(33) Bachmair, A.; Finley, D.; Varshavsky, A. In Vivo Half-Life of a Protein Is a Function of Its Amino-Terminal Residue. Science 1986, 234 (4773), 179–186. 10.1126/science.3018930.

(34) Ottoz, D. S. M.; Rudolf, F.; Stelling, J. Inducible, Tightly Regulated and Growth Condition-Independent Transcription Factor in Saccharomyces Cerevisiae. Nucleic Acids Res. 2014, 42 (17), e130. 10.1093/nar/gku616.

(35) Nguyen, A. W.; Daugherty, P. S. Evolutionary Optimization of Fluorescent Proteins for Intracellular FRET. Nat. Biotechnol. 2005, 23 (3), 355–360. 10.1038/nbt1066.

(36) Gietz, R. D.; Schiestl, R. H.; Willems, A. R.; Woods, R. A. Studies on the Transformation of Intact Yeast Cells by the LiAc/SS-DNA/PEG Procedure. Yeast Chichester Engl. 1995, 11 (4), 355–360. 10.1002/yea.320110408.

(37) Adams, P. D.; Afonine, P. V.; Bunkóczi, G.; Chen, V. B.; Davis, I. W.; Echols, N.; Headd, J. J.; Hung, L.-W.; Kapral, G. J.; Grosse-Kunstleve, R. W.; McCoy, A. J.; Moriarty, N. W.; Oeffner, R.; Read, R. J.; Richardson, D. C.; Richardson, J. S.; Terwilliger, T. C.; Zwart, P. H. PHENIX: A Comprehensive Python-Based System for Macromolecular Structure Solution. Acta Crystallogr. D Biol. Crystallogr. 2010, 66 (Pt 2), 213–221. 10.1107/S0907444909052925.

(38) McCoy, A. J.; Grosse-Kunstleve, R. W.; Adams, P. D.; Winn, M. D.; Storoni, L. C.; Read, R. J. Phaser Crystallographic Sohware. J. Appl. Crystallogr. 2007, 40 (Pt 4), 658–674. 10.1107/S0021889807021206.

(39) Emsley, P.; Cowtan, K. Coot: Model-Building Tools for Molecular Graphics. Acta Crystallogr. D Biol. Crystallogr. 2004, 60 (Pt 12 Pt 1), 2126–2132. 10.1107/S0907444904019158.

(40) Brünger, A. T. Free R Value: A Novel Statistical Quantity for Assessing the Accuracy of Crystal Structures. Nature 1992, 355 (6359), 472–475. 10.1038/355472a0.

