## Supplementary Figures for "Contributions of hyperactive mutations in M^pro^ from SARS-CoV-2 to drug resistance"

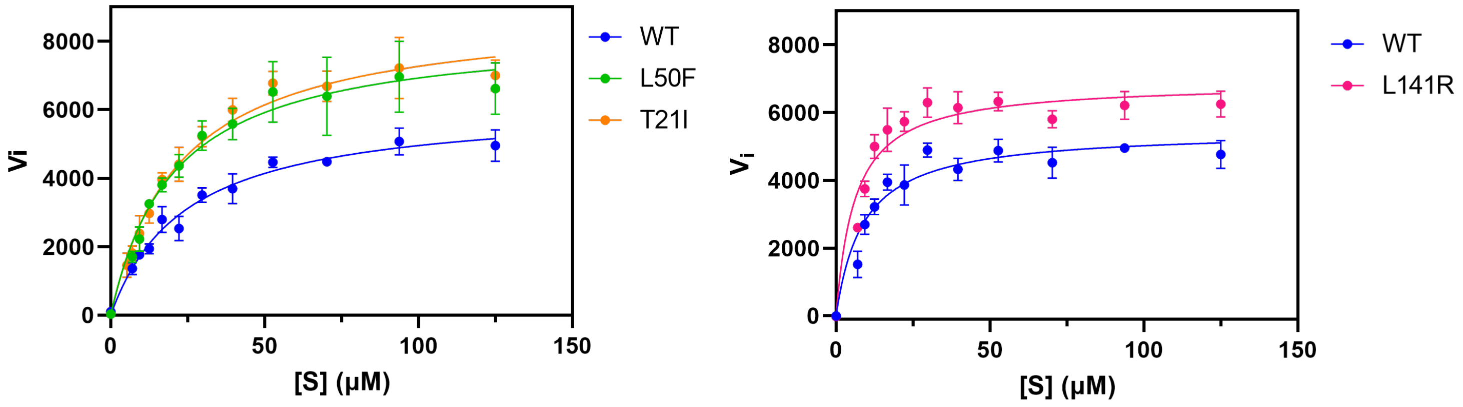


| $M^{pro}$ mutant | Relative $k_{cat}/K_m$ |
|---|---|
| **WT** | 1.0 |
| **T21I** | 1.6 |
| **L50F** | 1.7 |
| **L141R** | 1.8 |

Figure S1. T21I, L50F, and L141R $M^{pro}$ exhibit elevated enzymatic activity. Enzymatic activity was measured using a fluorogenic Nsp4/5 peptide substrate. L141R was performed using a different batch of peptide substrate and its relative activity was compared to WT measured with the same batch of peptide on the same day. Curves were fit using the Michaelis-Menten equation in Prism software. Each measurement was performed in triplicate.

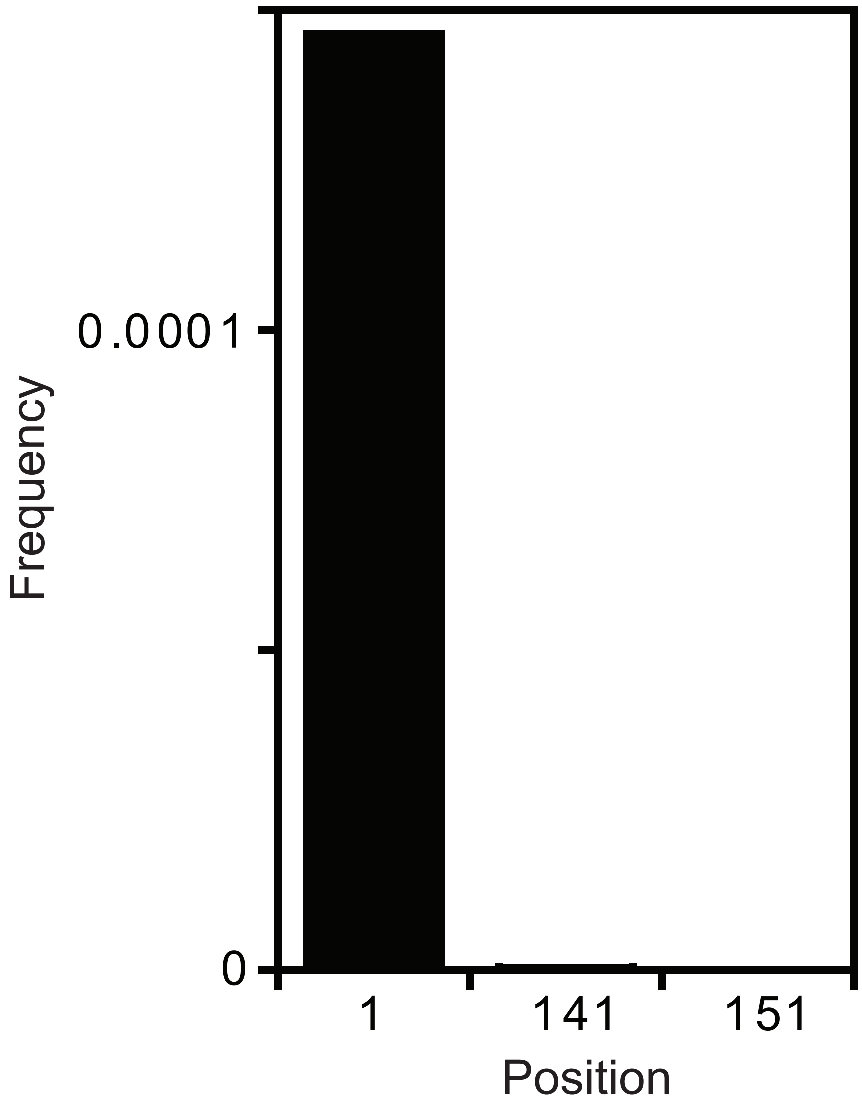


Figure ?2. Frequency of hyperactive mutations in sequenced isolates. Based on sequencing data reported as of September 7, 2023. Includes mutations identified as hyperactive in our yeast screen.

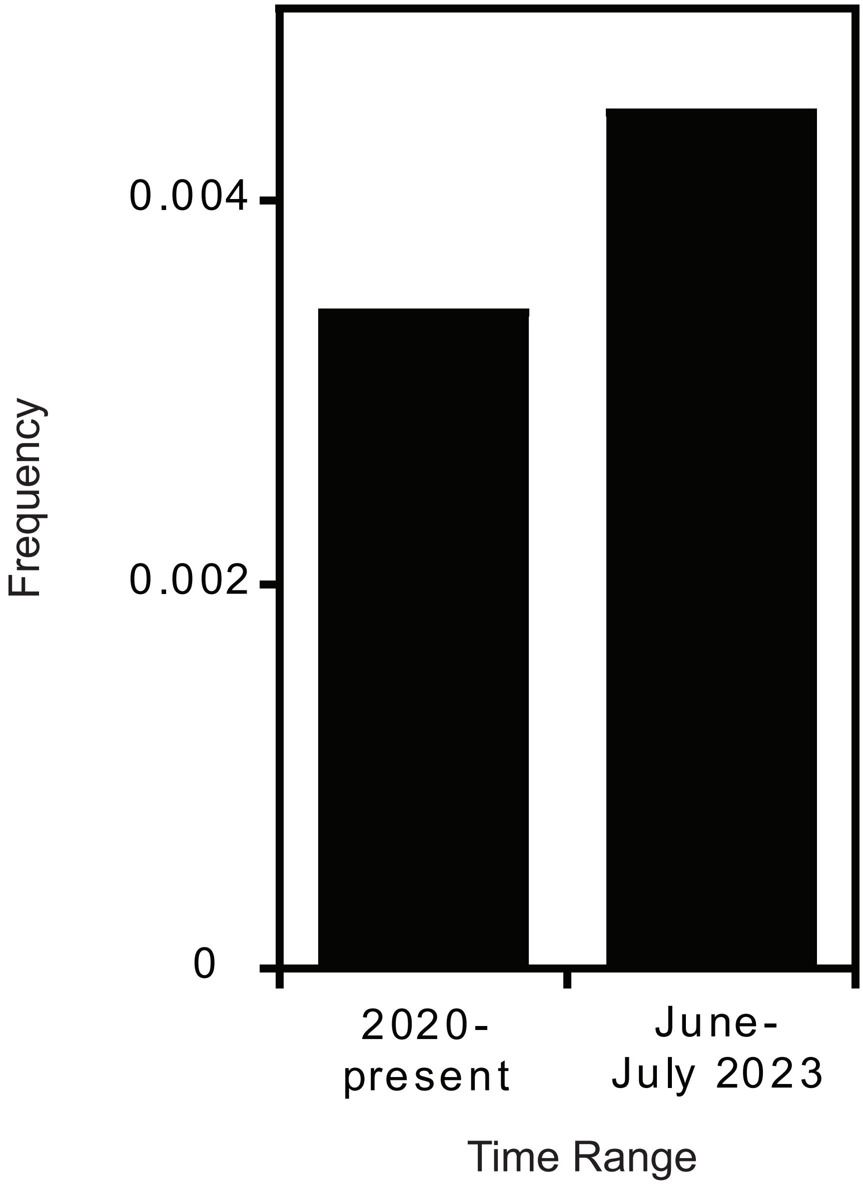


Figure S3. Frequency of mutations of resistance concner in sequenced SARS-CoV-2 isolates. Mutations with resistance and/or hyperactivity in yeast screens were tabulated and normalized to total sequenced isolates in each time window.

Table S3: Crystallization and refinement statistics for SARS-CoV-2 M[pro] mutations with small molecule inhibitors.

| **SC2-M[pro] Mutant-Inhibitor** | **L50F-PF-07321332** | **L141R-PF-00835231** | **N142P-PF-00835231** |
|---|---|---|---|
| PDB ID | 8E5C | 8DT9 | 8E4W |
| Data Collection | | | |
| Location | Home source | Home source | Home source |
| Resolution range (Å) | 25.41-2.2 (2.279-2.2) | 23.58-2.0 (2.071-2.0) | 23.75-2.75 (2.848-2.75) |
| Space group | $P\ 2_1\ 2_1\ 2$ | $P\ 1\ 2_1\ 1$ | $P\ 1\ 2_1\ 1$ |
| a, b, c (Å) | 45.1, 63.7, 105.3 | 54.5, 98.5, 58.5 | 55.1, 98.2, 58.8 |
| $\alpha$, β, γ (°) | 90, 90, 90 | 90, 107.4, 90 | 90, 107.8, 90 |
| Total reflections | 32011 (3082) | 174137 (12505) | 79795 (6496) |
| Unique reflections | 16012 (1546) | 39784 (3939) | 15527 (1525) |
| Multiplicity | 2.0 (2.0) | 4.4 (3.2) | 5.1 (4.3) |
| Completeness (%) | 99.82 (99.55) | 99.82 (99.60) | 99.79 (100.00) |
| Average $I/\sigma$ | 7.25 (1.95) | 14.51 (2.22) | 8.06 (2.10) |
| Wilson B-factor | 17.10 | 20.22 | 26.46 |
| $R_{merge}$[a] | 0.08596 (0.3643) | 0.07858 (0.4677) | 0.1813 (0.6675) |
| $CC_{1/2}$ | 0.988 (0.707) | 0.997 (0.8) | 0.977 (0.706) |
| Refinement | | | |
| $R_{factor}$[c] | 0.1955 (0.2310) | 0.1851 (0.2565) | 0.1879 (0.2309) |
| $R_{free}$[d] | 0.2538 (0.2767) | 0.2426 (0.3250) | 0.2410 (0.2572) |
| RMSD[b] in: | | | |
| Bond length (Å) | 0.004 | 0.004 | 0.005 |
| Bond angles (°) | 0.61 | 0.57 | 0.74 |
| Ramachandran: | | | |
| Favored (%) | 97.36 | 97.18 | 95.71 |
| Allowed (%) | 2.64 | 2.82 | 4.29 |
| Outliers (%) | 0.00 | 0.00 | 0.00 |
| Rotamer outliers (%) | 0.39 | 1.38 | 2.02 |
| B-factors | | | |
| Average | 17.67 | 24.69 | 25.89 |
| Macromolecules | 16.18 | 23.44 | 25.74 |
| Ligand | 24.21 | 30.99 | 26.19 |
| Solvent | 26.95 | 34.00 | 28.81 |

[a] $R_{sym} = \Sigma \mid I - \langle I \rangle \mid / \Sigma I$, where $I$ = observed intensity, $\langle I \rangle$ = average intensity over symmetry equivalent.
[b] RMSD, root mean square deviation.
[c] $R_{factor} = \Sigma \|\, F_o| - |F_c\| / \Sigma|F_o|$.
[d] $R_{free}$ was calculated from 5% of reflections, chosen randomly, which were omitted from the refinement process.
Statistics for the highest-resolution shell are shown in parentheses.
